# Controlled adhesion, membrane pinning and vesicle transport by Janus particles

**DOI:** 10.1101/2022.01.19.474912

**Authors:** Eleanor J. Ewins, Koohee Han, Bhuvnesh Bharti, Tom Robinson, Orlin D. Velev, Rumiana Dimova

## Abstract

The interactions between biomembranes and particles are key to many applications, but the lack of controllable model systems to study them limits the progress in their research. Here, we describe how Janus polystyrene microparticles, half coated with iron, can be partially engulfed by artificial cells, namely giant vesicles, with the goals to control and investigate their adhesion and degree of encapsulation. The interaction between the Janus particles and these model cell membrane systems is mediated by electrostatic charge, offering a further mode of modulation in addition to the iron patches. The ferromagnetic particle coatings also enable the ability to manipulate and transport the vesicles by magnetic fields.

---

Interactions of particles with biomembranes are widely studied due to their relevance in multiple current and potential applications, such as in medical imaging,^1^ or as antimicrobial agents,^2^ or to understand the negative environmental impact of microplastics.^3^ In order to take full advantage of these applications, it is important to understand the underlying mechanisms and parameters that govern the adhesion and engulfment of particles by membranes. In cells, this process is referred to as endo- or phagocytosis. Model membrane systems are commonly implemented for studying such processes.^4^ Among these systems are giant unilamellar vesicles (GUVs)^5^, which mimic the cell size and the curvature of the plasma membrane without the compositional complexity of live cells (which includes a wide variety of lipid species, proteins or the associated cytoskeleton).^6^ When investigating the parameters that dictate particle-membrane interactions, one vital aspect to consider is the role of the particle properties on the interaction potentials. For example, previous studies have examined how size, shape and surface chemistry impact the interactions of particles with cells^7^ or cellular mimetic systems^4k, 8^. What has yet to be explored experimentally is the behaviour of the endocytosis process for non-receptor mediated interactions between biomimetic membranes and synthetic particles with surface asymmetry.

By using micron-sized particles with two regions of distinctly different surface properties, commonly known as Janus particles, we investigate to what extent spatially varied surface properties govern the microsphere adhesion and engulfment by GUVs. Such anisotropic particles are of particular interest as they provide the opportunity to combine two different and sometimes incompatible properties within a single particle.^9^ They also provide means to quantify rotational dynamics due to their broken symmetry,^4j^ which could be a promising method for the further study of particle endocytosis^10^ or self-propelled guided transport and membrane deformation.^4j, 4k^ We find that by altering the particle surface chemistry in a spatially dependent manner, the adhesion and engulfment of microparticles becomes controllable. In addition to this, we make use of the iron oxide coating on the particle hemisphere to manipulate the particle-vesicle pairs using an external magnetic field gradient.

To select a GUV-particle combination exhibiting adhesion, we first performed a high-throughput screening assay with large unilamellar vesicles (LUVs) of different compositions. The LUVs were prepared via extrusion and incubated with particles of different surface chemistries (for details see Section S1.2 in the Supporting Information, SI). LUVs were composed of 1,2-dioleoyl-sn-glycero-3-phosphocholine (DOPC) with 40 mol% either 1,2-dioleoyl-sn-glycero-3-phospho-(1’-rac-glycerol) (DOPG - negative) or 1,2-dioleoyl-3-trime-thylammonium-propane (DOTAP - positive) to modulate the membrane charge; the lipid structures are given in Fig. S1. The microparticles used were polystyrene, either functionalised with sulphate or amine surface groups, resulting in negatively and positively charged surfaces respectively at neutral pH. The use of fluorescently labelled LUVs, containing 0.5 mol% 1,2-dipalmitoyl-sn-glycero-3-phosphoethanolamine-N-(lissamine rhodamine B sulfonyl) (Rh-DPPE) allowed for qualitative analysis of the relative affinity between the particle-vesicle combinations.

We observed a clear affinity of the positively charged LUVs to the negative polystyrene particles, see SI Fig. S3. Subsequently, we investigated the interactions between positively charged GUVs (containing DOTAP) and uniform and Janus particles exposing negative (sulphate) surface. The GUVs were prepared via electroformation (SI Section S1.2) from DOPC, doped with varying amounts of DOTAP (0-5 mol%; above this fraction the GUV quality was very poor and adhesion was not explored quantitatively) and a small amount of fluorescently labelled lipid (0.5 mol%Rh-DPPE) in 200 mM sucrose. Adhesion to neutral membranes (0% DOTAP) was not observed. The negative charge of the uniform polystyrene particles at neutral pH (here pH 7.45) can be attributed to the presence of sulphate groups on the surface.

Iron-patched Janus microspheres were prepared from the uniform polystyrene particles using metal vapour deposition technique (see SI Section S1.3)^11^ resulting in a hemispherical patch of 5 nm of chromium and 20 nm of iron. Note that the iron patch on the surface of microspheres transforms to iron oxide (Fe_2_O_3_) upon their resuspension in an aqueous environment.^11^ This region of the particle appears darker in brightfield images, see Fig. S2C.

Both the uniform and Janus particles were dispersed in hypertonic glucose solutions (see SI Section S1 for details); when the particles are incubated with the GUVs, this generates excess membrane area via osmotic deflation of the vesicles. We observed that this deflation step was necessary for particle engulfment to occur to any extent. Figure 1 shows example images of the two samples; 5% DOTAP GUVs that, typically, completely engulf the uniform, negatively charged polystyrene particles (Fig. 1A), whereas the Janus particles are partially engulfed (half-wrapped by the membrane) exhibiting pinning of the membrane contact line (Fig. 1B). For the Janus particles, the region of the particle in contact with the membrane is the polystyrene half (light region on particle in bright field images Fig. 1B) and the iron-patched half (dark region) remains at the periphery and restricts engulfment. The complete engulfment of the uniform polystyrene particles suggests that there is strong adhesion of the GUV membrane and the microsphere. This is corroborated by observations showing that the surface of Janus particles is only partially covered by LUVs, see Fig. S4.

**Fig. 1.**
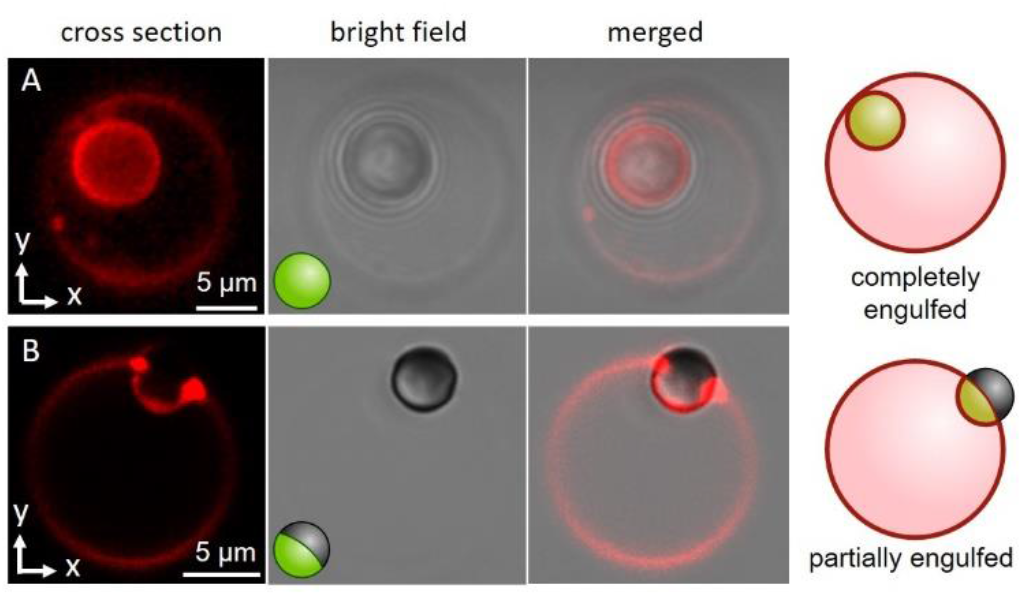
Confocal cross section and bright-field microscopy images of 5% DOTAP GUVs (red fluorescence) in contact with microparticles with uniform (A) and Janus (B) surface chemistries. (A) A 6 μm negatively charged (sulphate surface groups) polystyrene particle is completely engulfed by the GUV. (B) A 4 μm Janus particle, half negatively charged polystyrene (the same surface chemistry as in panel A) and half with a thin coating of iron, is partially engulfed by the GUV. The contact line of the adhered vesicle approximately corresponds to the iron-coated region of the particle surface, which can be seen in the brightfield image as the section of the particle that is darker (orientated away from the vesicle surface). The sketches illustrate how the particle surface is orientated in relation to the membrane and the state of engulfment.

These observations imply that the degree and energy of particle engulfment could be tuneable by altering the proportion of the particle surface that has a strong interaction with the lipid membrane, here shown on half-coated (Janus) or uniform particles. We further investigated this concept by measuring how the penetration depth of the particles into vesicles varied, both as a function of the particle surface (uniform or Janus) and membrane charge (GUVs doped with either 5 or 1% DOTAP). These results are displayed in Fig. 2, together with a schematic diagram illustrating how we define the penetration depth of the particle. Our definition of the penetration depth is comparable to that introduced by Dietrich et al.^12^, who quantitatively analysed the uptake of uniform particles by GUVs. The penetration depth is normalized which allows us to compare particles and vesicles of different sizes. The images shown in Fig. 1 have a “close-to-ideal” orientation of the vesicle-particle system, shown in a merged (brightfield and confocal fluorescence) image which directly reveals the penetration depth. However, in the entire sample, the particles can exhibit different positions with respect to the vesicle centre, and are typically located at the lower part of the GUV, making it nearly impossible to resolve the particle position from such projected images. We thus further develop the approach in Ref. ^12^ taking advantage of the improved resolvability provided by confocal microscopy, especially in the axial direction - see SI Section S3 for a detailed description of the penetration depth estimates from 3D confocal stacks.

**Fig. 2.**
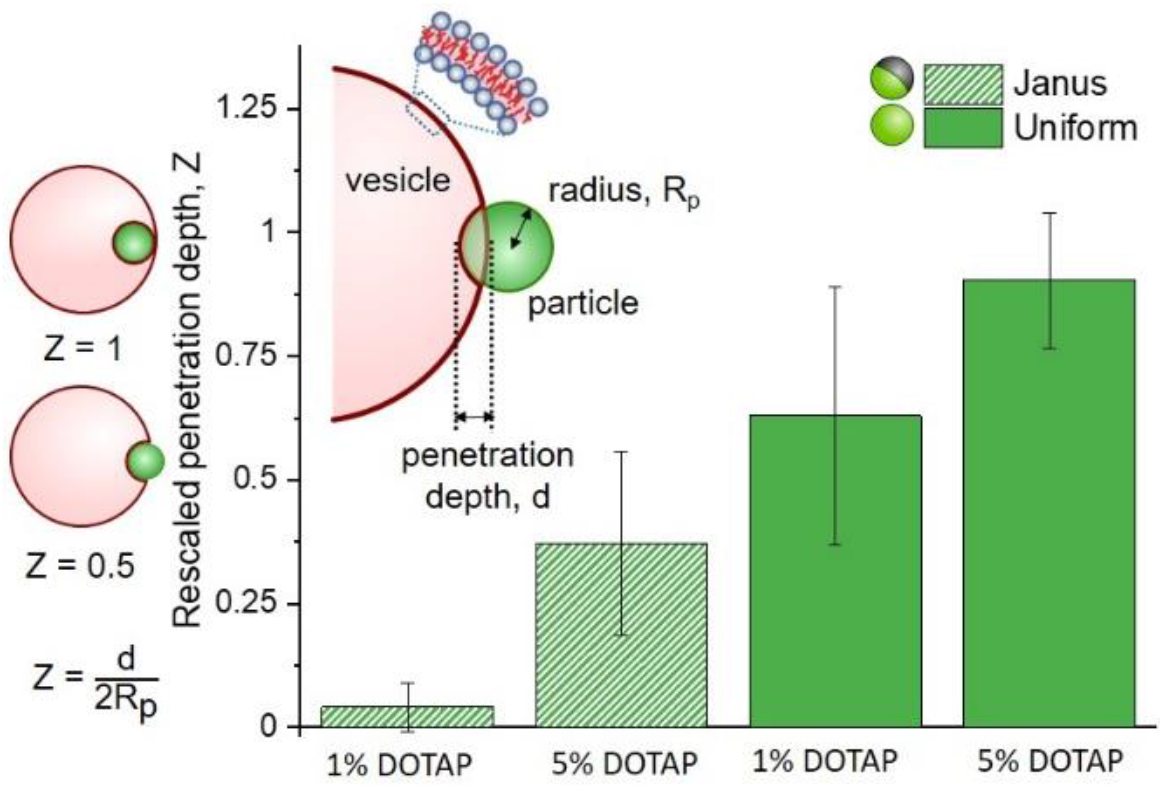
Penetration depths of Janus and homogeneous microparticles into GUVs composed of DOPC with 1 or 5% DOTAP (positively charged). Penetration depth d sketched in the inset is normalised by the particle diameter 2R_p_, so that different sized particles can be directly compared. The sketches on the left are representative of rescaled penetration depths of Z = 0.5 and Z = 1, of an arbitrary particle (green) into a vesicle (red). Janus particles, with half of their surface coated in metal, do not penetrate further than their radial depth into the vesicles (Z < 0.5), whereas homogeneous particles penetrate further. Between the two particle types, both particles penetrate further into membranes containing a larger proportion of positively charged lipid (an effective increase in adhesion energy). The analysis was carried out for the following number of vesicle-particle pairs: 1% DOTAP, Janus n = 8; 5% DOTAP, Janus n = 10; 1% DOTAP, uniform n = 7; 5% DOTAP, uniform n = 4. The errors are standard deviations. The vesicle diameters were in the range of 10-42 μm, and the particle diameters were 3.7 – 4.05 μm (Janus) and 5.8 – 6.3 μm (uniform).

The analysis of multiple interactions shows that uniform particles (solid bars, Fig. 2) penetrate further into the vesicles than Janus particles (hatched bars, Fig. 2), which we would expect on the basis of the qualitative observations in Fig. 1. The metallic regions supposedly repel the membrane or act as areas of the particles’ surfaces with lower adhesion energy, without a significant energy gain if the membrane would continue deforming to wet this part of the surface. Therefore, the wetting of the particle surface stops and the particles only partially penetrate into the vesicle. The contact line is pinned at the boundary between the polystyrene and iron oxide. Based on the data shown in Fig. 2, we can also see that there isn’t any single definitive penetration depth for each condition. This is most likely due to the challenges that arise from GUV and particle preparation: (i) For GUVs produced from a lipid mixture, it has been shown that the individual vesicle compositions vary^13^, so is the vesicle size relative to that of the particles. (ii) Vesicles with similar sizes can exhibit variable excess area for wrapping the particles (note that the preparation protocol does not allow for control over the initial vesicle area-to-volume ratio, and that the volume is osmotically fixed during particle engulfment). (iii) The surface chemistry of the polystyrene part of the Janus particles may differ from that of the homogeneous particles because of preparation steps (see SI section S1.3). (iv) Potential variation in the membrane spontaneous curvature could be expected (due to charge asymmetry^13^) which has been predicted to play a crucial role in particle engulfment^14^.

We also see that for both uniform and Janus particles the penetration depths have a dependence on the membrane charge (percentage of positively charged DOTAP, see Fig. 2); essentially, increasing the adhesion energy between the particle and the vesicle results in increased penetration of the particle into the vesicle. This process is seemingly governed by charge. To further investigate this, we examined the effects of increasing ionic strength, i.e. adding salt (150 mM NaCl) to the system, by observing the adhesion of LUVs doped with DOTAP to the 6 μm uniform microparticles.

Figure 3A shows images of particles with adhered LUVs in the presence and absence of salt (brightfield images showing the location of the particle can be found in Fig. S6). To assess the effect of salt, we quantified the fluorescence intensity of the particles; see Fig. S7A for an example along with an image of a bare, non-fluorescent particle (Fig. S7B, in the absence of LUVs). In 150 mM NaCl, which is expected to screen the charges, the intensity of the adhered LUVs is roughly 1.8 times lower than for the samples containing only sugars (0 mM NaCl), Fig. 3C; the scatter in fluorescence intensity values is possibly due to the small size variation between particles, as all particles are measured at the same distance from the glass surface. The results demonstrate that the interactions depend on electrostatics, in correlation with our observations of increased penetration depths in GUVs with a higher DOTAP content.

**Fig. 3.**
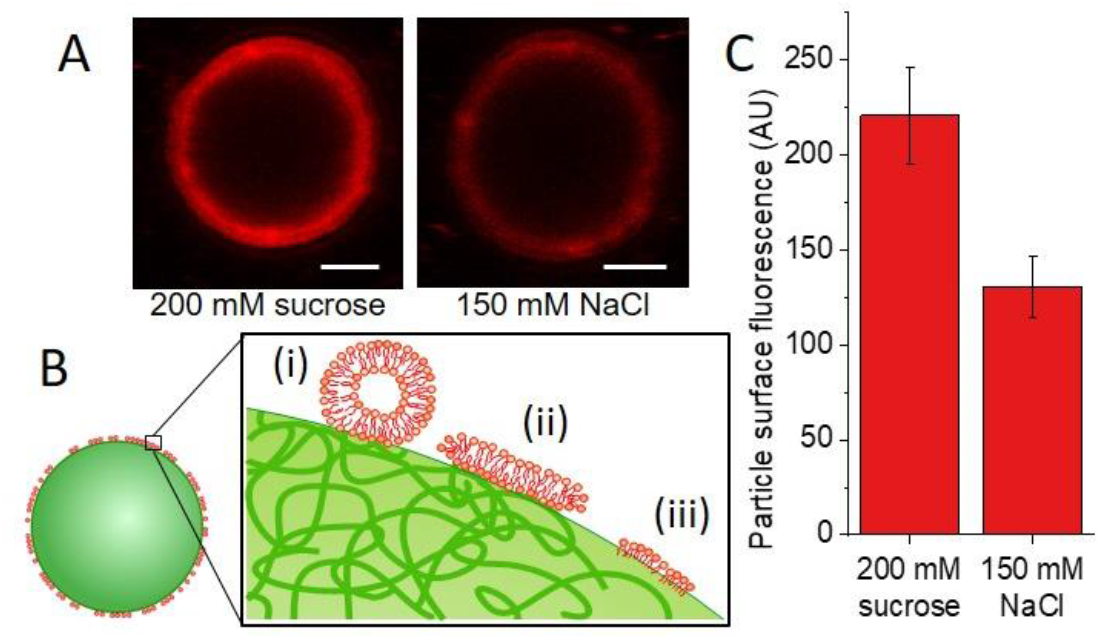
Effects of salt on the adhesion of positively charged membranes to negatively charged uniform particles. (A) LUVs (100 nm in size) composed of DOPC/DOTAP/Rh-DPPE 94.5/5/0.5 molar ratio, adhere to negatively charged polystyrene 6 μm particles to varying extents depending on the salt concentration in the solution, as deduced from confocal cross section images (scale bars: 2 μm). (B) The sketch (not to scale) represents possible reorganization of the red-labelled LUV membrane upon contact with the polystyrene particle surface (green) leading to non-uniform fluorescence over the particle surface: (i) adhesion, (ii) vesicle rupture and formation of a supported bilayer, or (iii) restructuring to a monolayer-like structure adhered to more hydrophobic patches on the particle surface. (C) In the presence of sugars (200 mM sucrose), the fluorescence intensity on the particle surface is higher by a factor of approximately 1.8; the presence of the salt ions partially screens the electrostatic interactions between the particles and the LUVs. The analysis is performed on 10 particles for each condition and error is the standard deviation of the mean fluorescence values.

However, these interactions do not appear to depend only on electrostatic attraction, as we still detect some fluorescence signal from LUVs adhered to the polystyrene microparticles in the presence of salt. Figure 3B includes a schematic diagram depicting different possible configurations of the LUV lipids and membrane adhering to the particle surface: (i) docked LUVs (single adhered LUVs appear to produce stronger signal; see comparison in Fig. S8); (ii) supported lipid bilayer (shown to be formed when LUVs adhere to silica particles and collapse^15^); and (iii) frustrated lipid monolayer adsorbed onto the hydrophobic regions of the latex surface as speculated by Dietrich et al.^12^

In addition to providing a region with a lower adhesion energy, the Fe_2_O_3_ patch on the Janus particles also provides magneto-responsiveness to the GUV-microparticle complex.^16^ In the presence of external magnetic field gradient, the Janus particles will move towards the regions of higher magnetic field intensity; this motion of particles is known as magnetophoresis. Such an approach is widely used with homogeneous magnetic particles in cell sorting protocols to isolate or enrich specific cell populations.^17^ Indeed, we observe magnetophoresis of the particle-vesicle complex of Janus particles and GUVs in the presence of a magnetic field (Fig. 4). Details for the setup for applying the magnetic field and imaging of vesicle displacement are given in the SI. GUVs were prepared from DOPC/DOTAP lipids in a 95/5 mol% ratio. When the source of the magnetic field is located to the lower left of the observation chamber, the particle and adhered vesicle move in this direction as they follow the field gradient (Fig. 4A). Conversely, when the source of the magnetic field is moved to the upper right corner of the chamber, the same particle-vesicle pair changes direction to move towards the magnetic field source (Fig. 4B). Throughout the observation, the particle remained adhered to the GUV, and it was possible to repeat similar manipulations with further particle-vesicle pairs, where the distance traversed was comparable to the size of the observation chamber (≈10 mm). Our approach of transporting vesicles by means of adsorbed Janus particles provides a facile and precise means of moving and sorting GUVs. The approach appears superior in terms of transporting the vesicles over large distances compared to mainly rotating them as shown with dipolar GUVs with charged domains.^18^

**Fig. 4.**
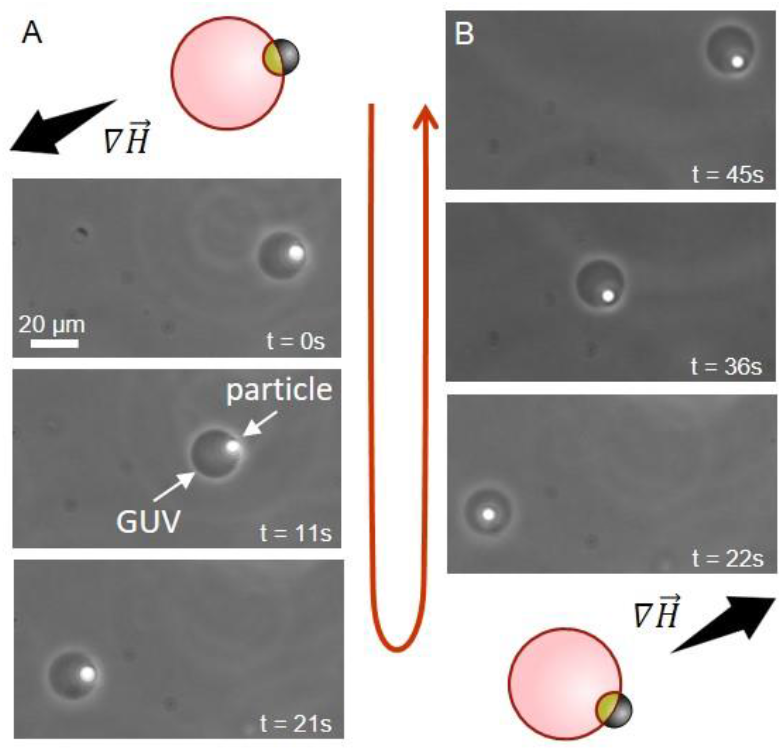
Time sequence demonstrating the transport of a GUV (dark circle) through the observation chamber via manipulation with an adhered Janus particle (white spot) in a non-uniform magnetic field; phase-contrast microscopy, see also SI Movie S1. The dense particle has sedimented to the lower half of the vesicle (out of focus), which is why in the transmitted light image it appears inside, but it is outside the GUV. The schematic diagrams indicate the initial particle-vesicle configuration and the magnetic field gradient they are exposed to, which causes the magnetophoresis of the particle. (A) The bar magnet is located to the bottom left of the chamber, causing the Janus particle to move towards the region with a higher field gradient, 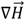. (B) The magnet is now located to the top right of the chamber and the Janus particle moves towards the region of increased field gradient. The particles are heavy (in the lower part of the GUV) and therefore having both vesicle and particle in focus is not always possible, which results in blurring of the image.

In conclusion, we have demonstrated the ability to control the extent of particle engulfment by biomembranes using Janus particles that have regions with different affinities for the GUV membranes. We also show that the particle iron oxide coating provides enhanced capabilities in terms of vesicle transport via magnetophoresis.

The degree of penetration varies as a combination of both particle surface asymmetry and lipid composition. This is coupled with a decrease in LUV adhesion in the presence of salt, which suggests that this system could be finely tuned to provide the desired degree of particle adhesion and penetration.

The use of Janus particles as a means to separate cell-like objects provides multiple opportunities for further development. The anisotropic surface could be used to limit the cells’ exposure to the damaging iron oxide,^19^ by creating regions of higher and lower affinity with the membrane. The exposed, non-binding region of the particle could also be functionalised so as to undergo exothermic surface reactions, as another means for generating self-propulsion.^20^ The selective adhesion demonstrated here could also be used as a template for spatially confined lipid sorting, as has been demonstrated by Liu et al.^18^ The controlled and directional force applied to GUVs could be used not only for sorting, but for quantitative characterization of the stiffness and moduli of the lipid membranes. This can be achieved both by the magnetic pull-off and torque. One could also consider a potential use of the entire vesicle-particle ensemble as a drug delivery system, with the manoeuvrability provided by the iron oxide coated particles, the lipids providing biocompatibility and the vesicles serving as a drug transporter.

## Supplementary Information

## List of abbreviations

AC: alternating current
BSA: bovine serum albumin
COM: centre of mass
DOPC: 1,2-dioleoyl-sn-glycero-3-phosphocholine
DOTAP: 1,2-dioleoyl-3-trimethylammonium-propane (chloride salt)
GUV: giant unilamellar vesicle
ITO: Indium tin oxide
LUV: large unilamellar vesicle
NA: numerical aperture
Rh-DPPE: 1,2-dipalmitoyl-sn-glycero-3-phosphoethanolamine-N-(lissamine rhodamine B sulfonyl) (Ammonium salt)
ROI: region of interest

## S1. Materials and methods

### S1.1. Materials

1,2-dioleoyl-sn-glycero-3-phosphocholine (DOPC), 1,2-dioleoyl-3-trimethylammonium-propane (chloride salt) (DOTAP) and 1,2-dipalmitoyl-sn-glycero-3-phosphoethanolamine-N-(lissamine rhodamine B sulfonyl) (Ammonium salt) (Rh-DPPE) were acquired from Avanti Polar Lipids (Alabaster, AL); the lipid structures are given in Fig. S1. 100 nm pore diameter polycarbonate membranes were obtained from Whatman (Maidstone, UK). Indium tin oxide (ITO) coated glasses (ITO film thickness < 100 nm, resistance 50 Ω) were obtained from Präzisions Glas & Optik (Iserlohn, Germany). Glucose, sucrose, sodium chloride and bovine serum albumin (BSA) were all obtained from Sigma Aldrich (Darmstadt, Germany). The polystyrene particles used for direct adhesion to LUVs and GUVs (Polybead^®^ microspheres, 6 μm non-functionalised polystyrene (exposed sulphate groups), 4 μm functionalised and non-functionalised (exposed amine or sulphate surface groups respectively)) were obtained from Polysciences (Germany). The uniform polystyrene particles used for Janus particle preparation were purchased from Bangs Lab Inc. (Indiana, USA). Iron and chromium pellets were purchased from Kurt J. Lesker Co. (Clairton, PA, USA).

**Fig. S1.**
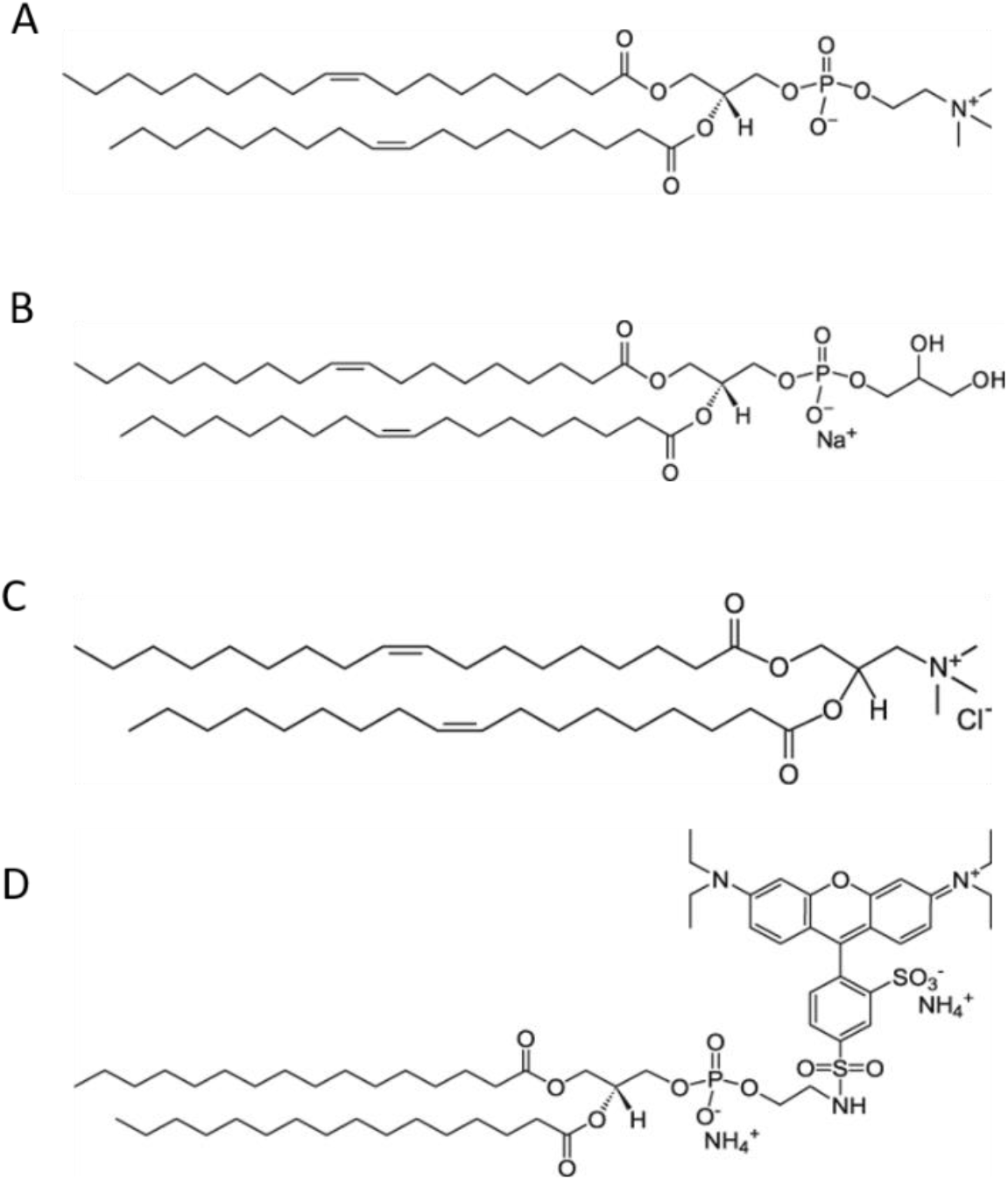
Structures of lipids used in this study. (A) 1,2-dioleoyl-sn-glycero-3-phosphocholine (DOPC). (B) 1,2-dioleoyl-sn-glycero-3-phospho-(1’-rac-glycerol) (sodium salt) (DOPG). (C) 1,2-dioleoyl-3-trimethylammonium-propane (chloride salt) (DOTAP). (D) 1,2-dipalmitoyl-sn-glycero-3-phosphoethanolamine-N-(lissamine rhodamine B sulfonyl) (ammonium salt) (Rh-DPPE). *Images from Avanti Polar Lipids*.

### S1.2. Vesicle preparation and mixing with particles

Large unilamellar vesicles (LUVs) were prepared via extrusion at room temperature.^1^ The lipid solution was prepared to 2.2 mL in chloroform at a concentration of 2 mM and composed of either DOPC/DOTAP/Rh-DPPE (59.5/40/0.5 molar ratio), DOPC/DOPG/Rh-DPPE (59.5/40/0.5 molar ratio) or DOPC/DOTAP/Rh-DPPE (94.5/5/0.5 molar ratio) and deposited in a small glass vial. The chloroform was evaporated first under a stream of N_2_ and then further dried under vacuum for 2 hours. The lipid film was then rehydrated with either 0.2 M sucrose or 150 mM NaCl to a final lipid concentration of 1.2 mM. The solution was vortexed for 2-5 minutes, obtaining multilamellar vesicles. The vesicle solution was then subjected to 11 cycles of extrusion through a 100 nm pore diameter polycarbonate membrane. For the adhesion studies, LUVs were mixed with the microspheres and incubated for 1 hour before imaging, in a vertical rotating mixer. The particles had a final concentration of 8.4 × 10^4^ particles/mL and the LUVs had a final lipid concentration of 0.83 mg/mL.

Giant unilamellar vesicles (GUVs) were prepared via the established electroformation protocol.^2^ Lipid solutions were prepared in chloroform at 4 mM with varying ratios of DOPC and DOTAP, as indicated throughout the text. Unless explicitly stated in the text, lipid solutions also contained 0.5 mol% Rh-DPPE fluorescent dye. A total volume of 16 μL of the lipid solution in chloroform was spread on two conductive ITO-coated glasses and dried under vacuum for 2 to 2.5 hours at room temperature. The ITO glasses, together with a rectangular Teflon spacer, were then assembled to form a chamber of 2 mL volume which was filled with 0.2 M sucrose. The chamber was then connected to a function generator which was used to apply an AC field (1.2 V, 10 Hz) for 1.5 hours at room temperature (for the lipid compositions containing dyes, the electroformation was performed in the dark). The GUVs were then removed from the growth chamber via pipetting and diluted 1:1 in a 0.21 M glucose solution (unless otherwise stated in the main text) containing dispersed particles. GUV suspensions and glucose solutions were measured and the osmolarity adjusted (glucose only) using an osmometer (Osmomat 030, Gonotec, Germany) such that the particle solution had higher osmolarity by ∼10 mOsm. The GUVs were incubated with the particles (final concentration of 8.4 × 10^3^ particles/mL) for 1 hour in a vertical rotating mixer before observation.

### S1.3. Janus particle preparation

Iron-patched Janus microspheres were prepared from the uniform polystyrene particles using a metal vapour deposition technique,^3, 4^ see sketch in Fig. S2A. Briefly, the polystyrene particles were concentrated and washed by centrifuging at 1500 ×*g* for 5 min and replacing the supernatant with MQ water; this was repeated 2-3 times. A convective assembly method was used to deposit particle monolayers on pre-cleaned glass slides.^5^ The dried particle monolayers were coated with hemispherical patches of chromium followed by a layer of iron (5 nm and 20 nm respectively) in a metal evaporator (Cooke Vacuum Products, model CV302). The thickness of the evaporated metals was monitored using a Maxtek Inc. TM350 thickness monitor equipped with SC-101 sensor crystals. The particles were then gently scraped from the deposition surface and resuspended in Milli-Q water. A SEM image of Janus particles is shown in Fig. S2B. The microparticles were then washed 3 times in Milli-Q water, via repeated centrifugation and removal of supernatant, before use. Note that the iron patch on the surface of microspheres transforms to iron oxide (Fe_2_O_3_) upon their resuspension in an aqueous environment.^3^ In some brightfield images (depending on the particle orientation), it is also possible to distinguish a region of the particle surface that is darker, which corresponds to the iron oxide cap, see Fig. S2C.

**Fig. S2.**
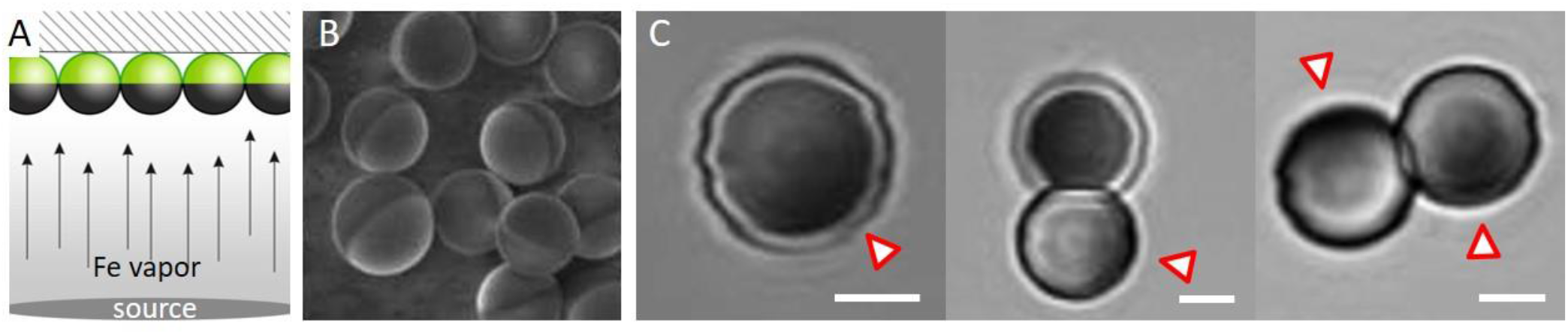
(A) Schematic of metal vapour deposition on a monolayer of polystyrene colloid spheres dried on a solid substrate, adapted from ^4^. (B) SEM image of 4 μm Janus particles adapted from^3^, clearly showing the two different surfaces on the same particle. (C) Brightfield images of 4 μm Janus particles in aqueous solution with visible darker patches (arrowheads pointing to them), corresponding to the iron oxide coating. Scale bars: 2 μm. The apparent distorted surface of the particle is an artefact of the slow scanning speed of the confocal microscope with which the images were acquired.

### S1.4. Imaging

Confocal imaging was performed on either a Leica confocal SP8 or SP5 setup (Mannheim, Germany). Rh-DPPE was excited with a 561 nm laser and the emission signal collected between 580-670 nm. The images were acquired with a 63x (1.2 NA) water immersion objective and 1 Airy unit. The subsequent image analysis is described in detail in section S2. Phase contrast imaging was performed on an Axio Observer D1 (Zeiss, Germany) microscope, equipped with a Ph2 20x (NA 0.5) objective and an ORCA R2 CCD camera (Hamamatsu, Japan).

### S1.5. Magnetic manipulation of Janus particles

A handheld bar magnet was used to generate a magnetic field gradient by placing the magnet close (approximately 2 cm from pole of magnet to glass) to the observation chamber. The magnet is formed from multiple blocks of Neodymium (dimensions 2.0 cm × 2.0 cm × 10.4 cm), with an approximate total magnetic field of 1 mT (at the magnet surface). To change the direction of the magnetic field, the position of the magnet was rotated approximately 180° around the observation chamber. During image acquisition the magnet was held in a fixed position.

## S2. Particle-membrane affinity with homogeneous and Janus particles

**Fig. S3.**
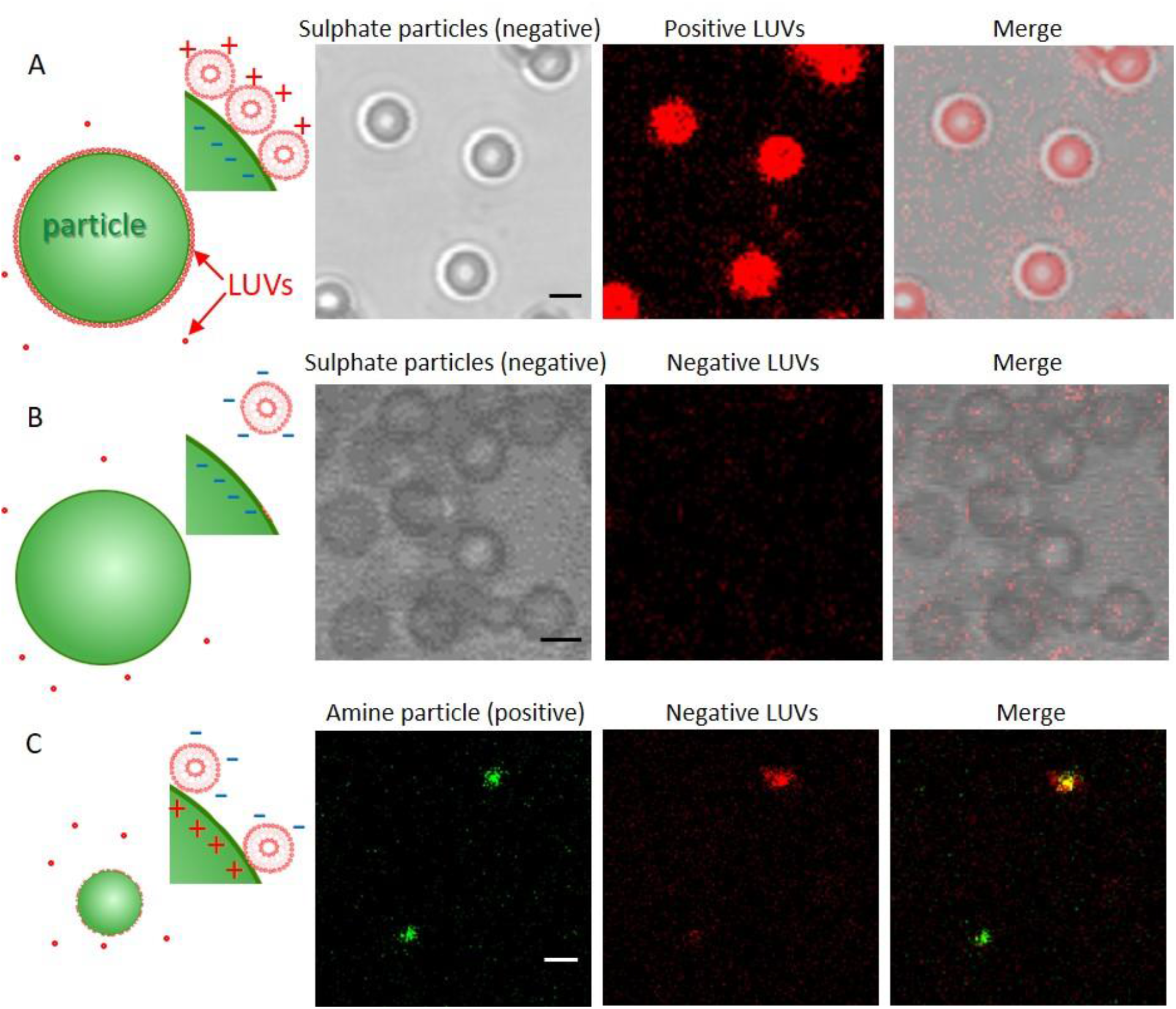
Adhesion of fluorescently labelled LUVs of varying lipid compositions (and labelled with 0.5 mol% Rh-DPPE, red) to microparticles with different surface charges shown with schematic representations of the different degrees of interactions and bright-field and confocal images. The sketches roughly represent the relatively large size of the microparticles (6 μm or 1 μm) relative to that of the LUVs (100 nm). (A) 6 μm polystyrene particles with a negative surface charge (sulphate groups) and positively charged LUVs (DOPC/DOTAP 60/40 mol%). The LUVs (as observed from the fluorescence signal from the red dye Rh-DPPE in the membrane) completely cover the surface of all of the particles in the sample. (B) The same 6 μm polystyrene particles and negatively charged LUVs (DOPC/DOPG 60/40 mol%). The LUVs do not adhere to the particles’ surface, which we conclude from the lack of fluorescence signal in the particles’ location in the merged image. (C) 1 μm polystyrene particles with amine functional groups (positively charged, labelled with green fluorescent dye) incubated with negatively charged LUVs (DOPC/DOPG 60/40 mol%, red) showed heterogeneous adhesion of LUVs to the particles’ surfaces, as can be seen from the difference in fluorescence signals from the red LUVs on the two particles (middle image). All scale bars correspond to 5 μm. The merged images on the right show overlay of the signal detected in the channels showing the particles and the LUVs individually.

**Fig. S4.**
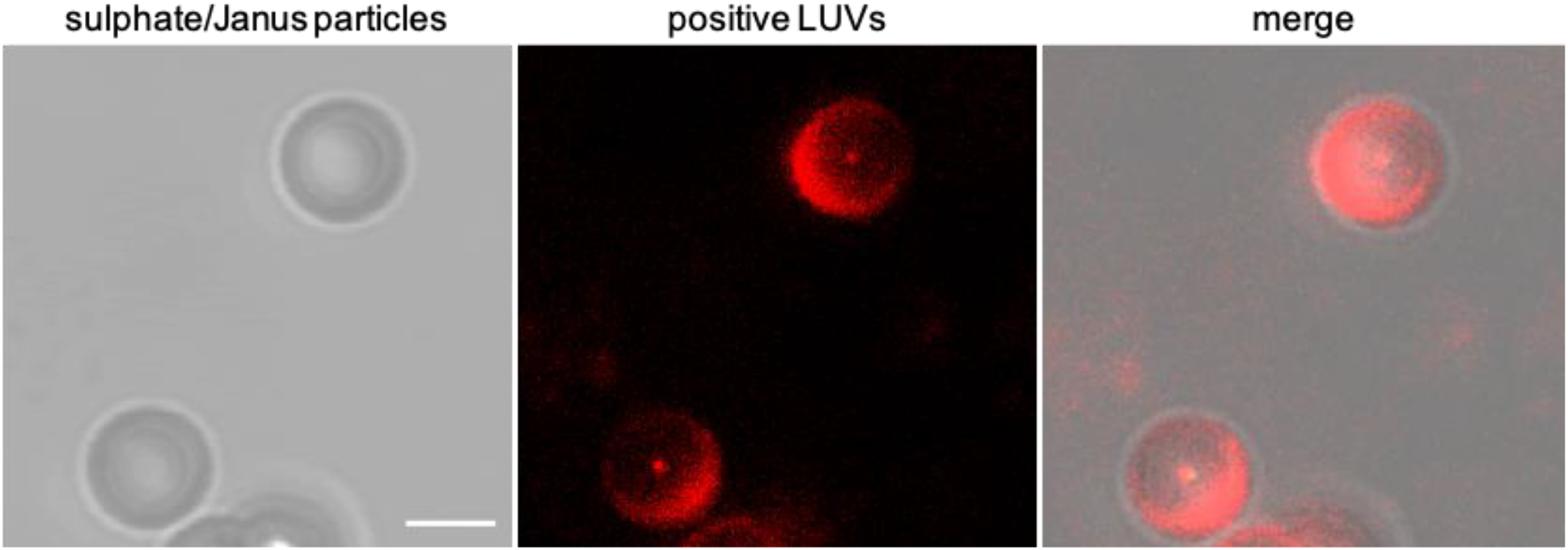
Preferential adhesion of LUVs to one region of Janus particles. DOTAP (positively charged) doped LUVs (labelled with Rh-DPPE visualized in red) only adhere to portions of Janus particle surfaces, indicating a stronger adhesion for one region compared to another. Scale bar: 3 μm. The non-adhesive region, exhibiting less fluorescence from the LUVs (middle image), corresponds to the metal patch, which appears darker in the brightfield image (left). The merged image shows an overlay of both.

## S3. Calculating particle penetration depth

The particle penetration depth into the GUVs was calculated from a stack of confocal z-slices of each vesicle-particle pair. First, the (x,y) centre of mass (*COM*_*V*_ in Fig. S5) of the vesicle was determined by fitting a circle to the vesicle contour in the z-stack with the largest diameter (the error on this value is the standard deviation of three such measurements on the same vesicle). The z-position of the COM(s) were calculated as the image number in the stack (e.g., z = 14) multiplied by the z step height (the error on this value was determined to be the z step height of the confocal stack, and is introduced when selecting the correct contour to measure). The centre of mass of the particle (*COM*_*P*_) was determined in the same way from the brightfield channel. The distance between the vesicle COM and particle COM was calculated using Equation 1:

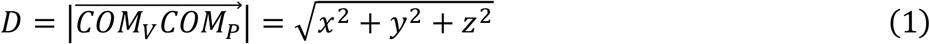

in which *COM*_*V*_ = [*x*_*V*_, *y*_*V*_, *z*_*V*_], *COM*_*P*_ = [*x*_*P*_, *y*_*P*_, *z*_*P*_] and subsequently *x* = *x*_*V*_ − *x*_*P*_ (and similarly for *y* and *z*). The depth of particle penetration into the vesicle (*d*) is defined as the distance between the vesicle membrane on the particle surface (solid line in contact with particle) and where it is projected to be (dashed red line) if the particle were not present (by assuming the vesicle is spherical). This distance is therefore calculated using the following equation:

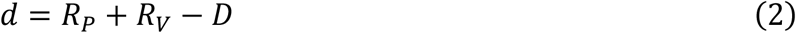

where *R*_*P*_ and *R*_*V*_ are the particle and vesicle radii respectively, as indicated in the diagram below.

**Fig. S5.**
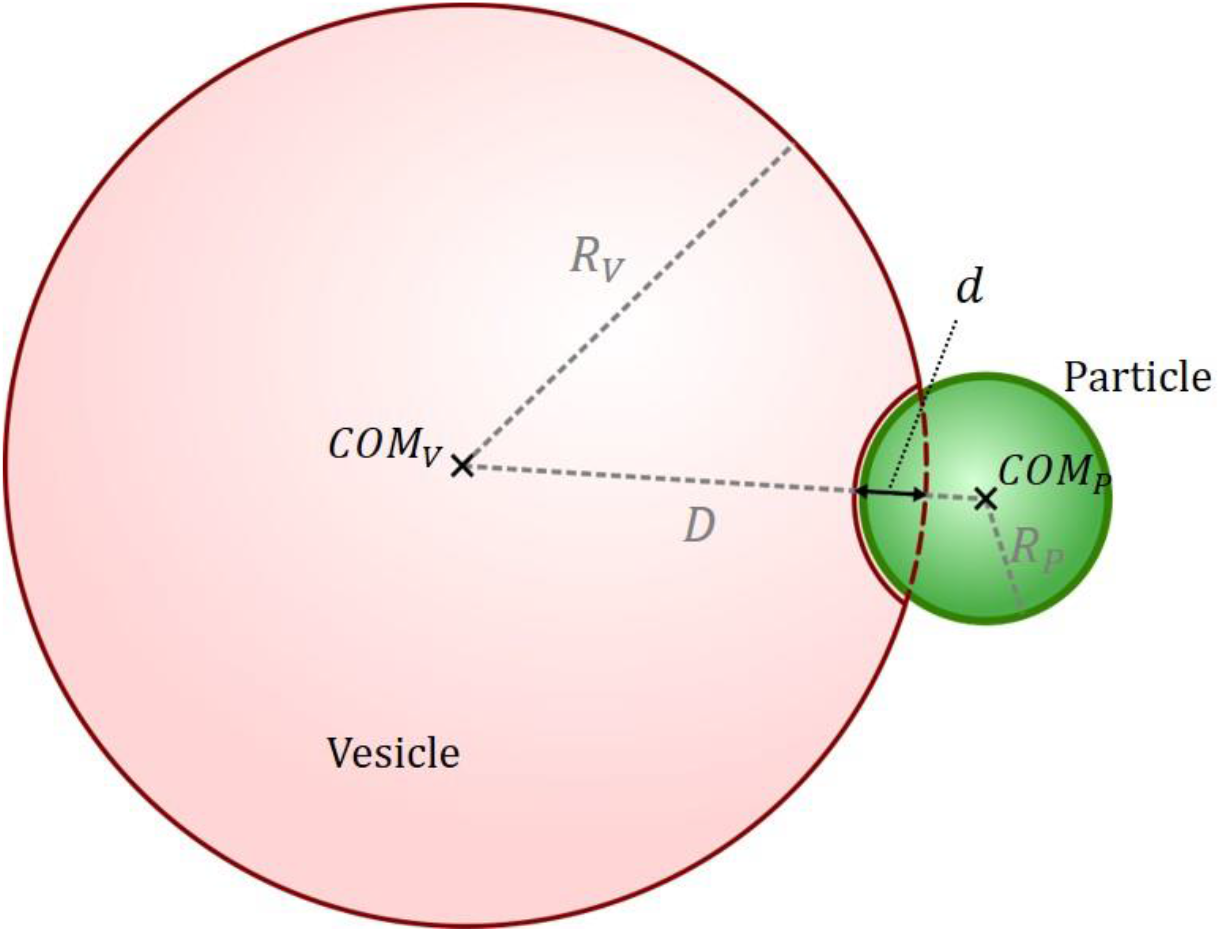
Schematic presentation of the vesicle and particle with indicated relevant dimensions of the system.

## S4. Imaging and evaluating LUV fluorescence on particle surface

**Fig. S6.**
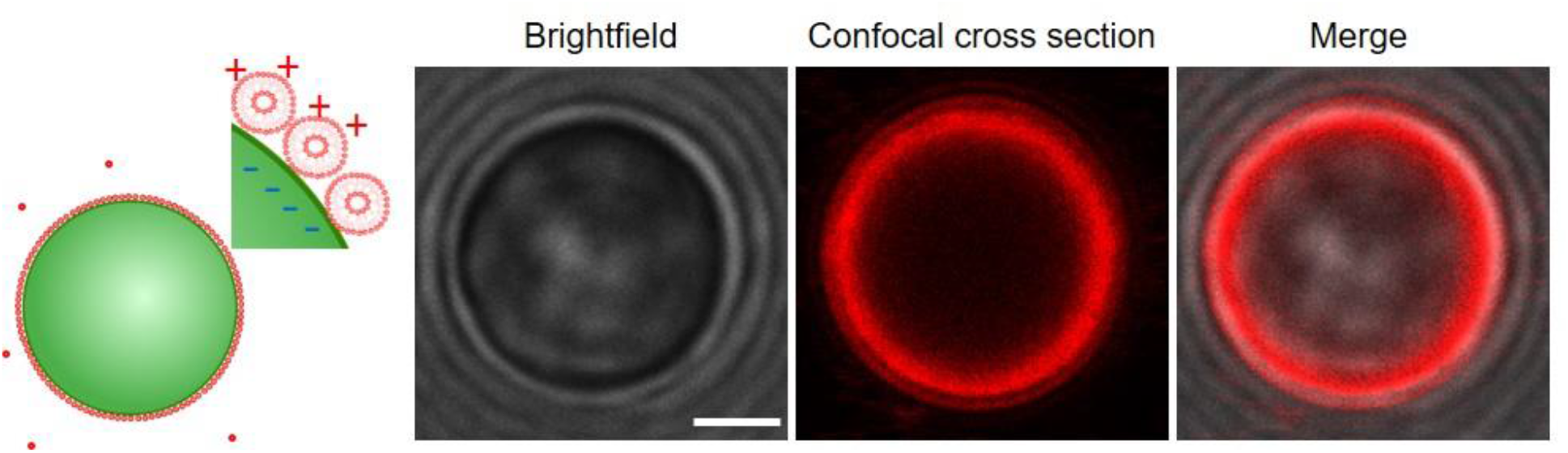
Brightfield and confocal cross-section with LUVs (red) adsorbed to the surface of a 6 μm polystyrene (sulphate surface groups, negative charge) particle (visible in brightfield image and overlay of the brightfield and red fluorescent channels) in the absence of salt. Scale bar 2 μm.

**Fig. S7.**
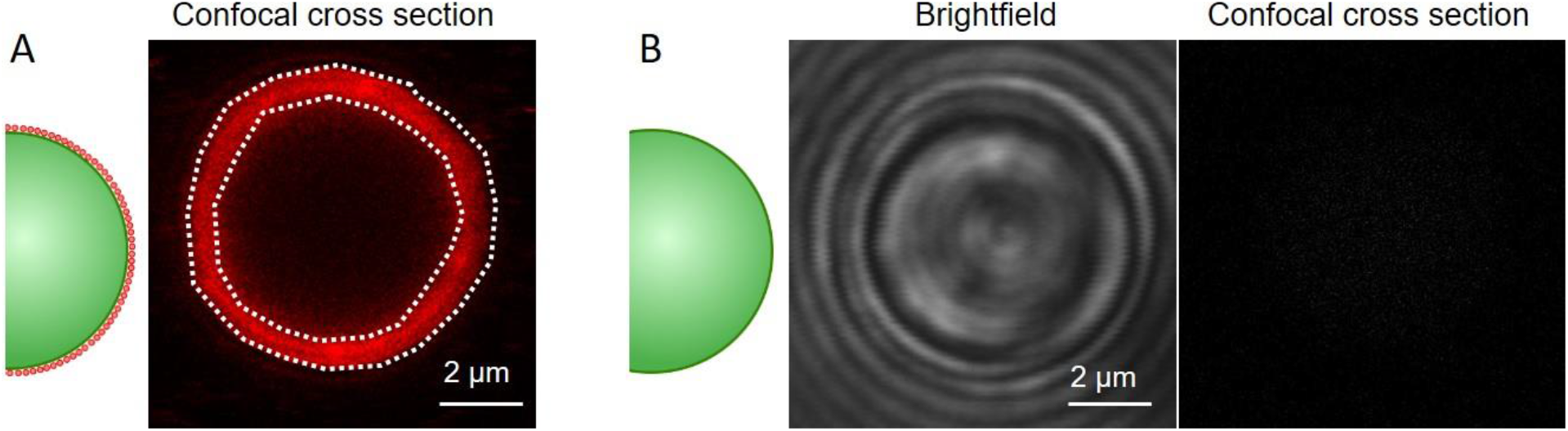
(A) Evaluating the fluorescently labelled LUV coverage on 6 μm polystyrene particles: a confocal cross section of one particle with dashed line indicating the ROI (region of interest) that is selected for fluorescence intensity analysis of LUV coverage. The analysis is carried out in LAS X, a confocal imaging and analysis software from Leica. As the particles are roughly of the same size, the equatorial cross-section is imaged at the same height from the glass for all particles. When the ROI is selected, regions with visible lipid/vesicle aggregation (and thus higher intensity) are excluded from the selection. Example demonstrated here is for 200 mM sucrose (no salt). (B) Images taken at the same imaging settings and image brightness/contrast for comparison of particles in the absence of LUVs demonstrate that none of the fluorescence signal results from particle reflection or autofluorescence (there is no red signal in the confocal panel).

**Fig. S8.**
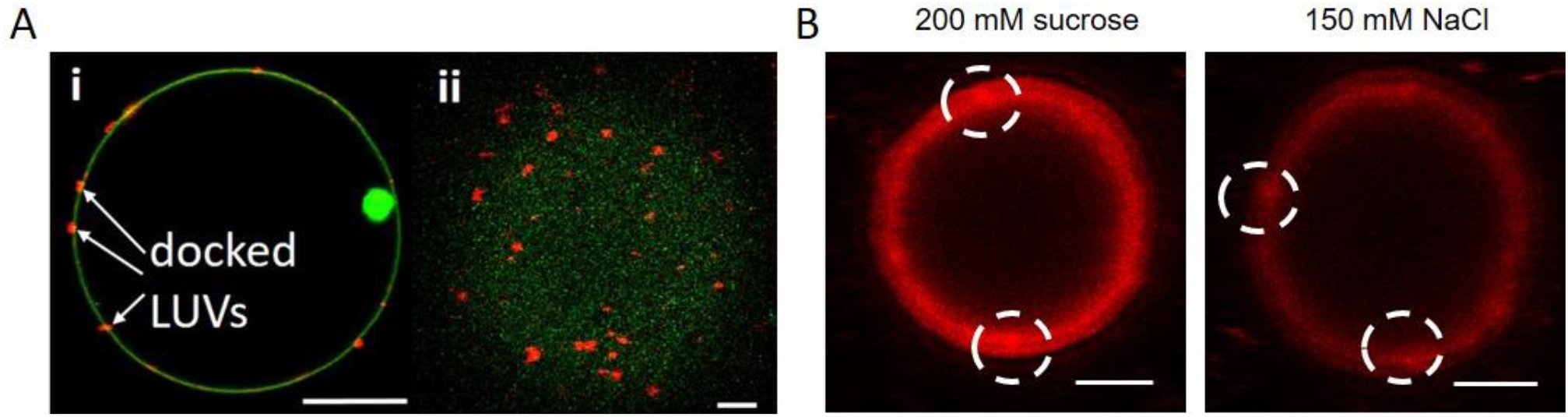
Comparison of LUVs fluorescence observed on GUV surface and on particle surface. (A) Confocal images of LUVs docked on the surface of a GUV; (i) mid-plane confocal cross section of GUV (green) with docked LUVs (red) indicated and (ii) upper GUV surface. Images adapted from ^6^. Scale bars (i) 10 μm, (ii) 2 μm. (B) Confocal image of LUVs present on the surface of a polystyrene particle in the conditions of Fig. 3 in the main text. Scale bars 2 μm. The fluorescence from individual docked LUVs (A) appear to produce stronger signal compared to the more homogeneous fluorescence that we observe over the particles (B), but certain spots of bright fluorescence (encircled in B) appear to result from LUV docking.

**Movie S1** Time series showing Janus particle adhered to a GUV moving through observation chamber in the presence of a magnetic field gradient; for details see Fig. 4 in the main text.

